# Intermolecular interactions of CYP3A enzymes and cytosolic sulfotransferases (SULT1E1, SULT2A1)

**DOI:** 10.64898/2026.10.14.759113

**Authors:** Anastasiya Tumilovich, Evgeniy Yablokov, Oksana Gnedenko, Diana Mednaia, Vadim Selin, Yuri Mezentsev, Pavel Ershov, Leonid Kaluzhskiy, Tatsiana Tsybruk, Irina Grabovec, Polina Shabunya, Andrei Gilep, Alexis Ivanov

## Abstract

Cytochrome P450 enzymes of the CYP3A subfamily (CYP3A4 and CYP3A5) and cytosolic sulfotransferases (SULT1E1 and SULT2A1) play key roles in the sequential oxidative and conjugative metabolism of steroid hormones and xenobiotics. While their coordinated transcriptional regulation and co-expression patterns are well documented, direct physical interactions between these phase I and phase II enzymes have not previously been demonstrated. In this study, we combined systems biology analysis, SPR biosensing, activity assays, and AlphaFold2 modeling to investigate PPIs between CYP3A4/3A5 and SULT1E1/2A1. STRINGdb and transcriptomic data suggested potential associations, but direct evidence was lacking. SPR confirmed high-affinity binding (Kd 0.29–1.8 nM); the SULT2A1–CYP3A5 pair gave 0.29 nM, while SULT2A1–CYP3A4 gave 1.8 nM. Notably, the presence of the sulfo group donor SULT PAPS, had minimal effect on binding affinity. Functional assays demonstrated that SULTs slightly inhibited CYP3A4 activity while moderately enhancing CYP3A5 activity in the presence of SULT1E1, both with and without PAPS, whereas the addition of CYPs leads to selectively increase in SULT2A1 activity in vitro by 38% (CYP3A4) and 108% (CYP3A5), without affecting SULT1E1. AlphaFold2 models revealed stable interfaces (715–1856 Å²) with 10–16 H-bonds and salt bridges. The results of our *in silico* tunnel analysis suggested a plausible alignment of substrate exit channels from CYPs with entry pathways to SULTs, supporting a mechanistic basis for substrate channeling in CYP–SULT complexes. These results indicate that the enzymes may form functional multienzyme complexes, offering new insights into coordinated steroid/xenobiotic metabolism and drug clearance.

**Significance statement:** This study provides the first direct evidence of high-affinity physical interactions between CYP3A and SULT enzymes, challenging the conventional view of independent metabolic phases. Our findings reveal that these interactions can reciprocally modulate enzymatic activities and suggest a structural basis for substrate channeling, offering a paradigm-shifting perspective on coordinated steroid and drug clearance that may reshape our understanding of hormone homeostasis and pharmacokinetics.

## 1. Introduction

Despite extensive studies of the individual catalytic properties of cytosolic sulfotransferases (SULTs) and microsomal cytochromes P450 (CYPs), the question of their functional coordination within the unified metabolic space of the cell remains open. Of particular interest are the CYP and SULT isoforms that exhibit overlapping substrate specificity toward key steroid hormones and physiologically important compounds (testosterone, dehydroepiandrosterone (DHEA, androstenolone), estradiol (E2), estrone (E1), cholesterol). Such SULTs include estrogen sulfotransferase (SULT1E1) and hydroxysteroid sulfotransferase (SULT2A1), while the associated CYPs are CYP3A4 and CYP3A5.^1–8^ Differences in the physicochemical properties of products derived from a common substrate upon action of CYPs or SULTs may represent one of the regulatory points of the metabolic and storage system for physiologically active steroid compounds in the human body.

The selection of these particular enzyme pairs as objects of the present study is due to the functional coupling of SULT1E1 and SULT2A1 with CYP3A4 and CYP3A5, which is not limited to competition for a common substrate. Sulfation products, long regarded exclusively as inactive metabolites, are now recognized not only as a circulating pool of inactive hormones but also as independent signaling molecules.^9,10^ It has been shown that steroid sulfates, such as estrone sulfate and DHEA sulfate, can directly modulate the activity of membrane receptors (GABA_A, NMDA) and transporters, acting as neuroactive steroids. Moreover, there is evidence for weak but dose-dependent transactivation of nuclear receptors (ER, AR) by sulfated forms.^11^ Consequently, the balance between SULT and CYP activity determines not only the metabolic clearance of the hormone but also the switching of its pharmacological and signaling profile — from a lipophilic agonist to a charged allosteric modulator.

Data from cellular models and transcriptomic analyses indicate that SULT2A1 and CYP3A4/3A5 can be co-expressed in the liver and intestine, promoting sequential metabolic transformation of steroids and xenobiotics: initial oxidation (phase I), followed by conjugation (phase II).^12^ By contrast, SULT1E1 displays a distinct tissue expression pattern and regulatory profile, enabling selective control of active estrogen levels.^13^

Additionally, results of analysis of normal and cancer tissue panels suggest the existence of shared regulatory networks among SULT and CYP genes, including positive correlations between SULT2A1 and CYP17A1 or CYP21A2 expression, further supporting their participation in an integrated endocrine-metabolic framework.^14^ Such co-regulation and co-expression are highly relevant to the pharmacokinetics of steroid hormones and drugs. Induction of CYP3A4/5 and SULT2A1 by PXR ligands can accelerate the clearance of both endogenous steroids and drugs, whereas repression of SULT1E1 may lead to elevated active estradiol concentrations, potentially influencing hormone-dependent pathologies.^15^

A panoramic bibliometric analysis of a set of 77 relevant articles collected from PubMed and Scilit databases for the period 2004–2025 revealed the presence of term co-occurrence of at least 12 CYPs (CYP1A1, 1A2, 1B1, 2B6, 2D6, 2C9, 2C19, 2E1, 3A4, 3A5, 7A1, and 8B1) and 5 SULTs (SULT1A1, 1A2, 1B1, 2A1, and 2B1) in the article’s texts (Figure S1 and S2, Supplementary material). Furthermore, the literature landscape analysis of a set of relevant articles revealed at least 5 characteristic types of functional associations between SULTs and non-steroidogenic microsomal CYPs. Table S1 (Supplementary material) demonstrates some examples of these associations: 1) ‘gene/protein co-expression under disease conditions’ [10.1016/j.xphs.2022.07.012, 10.1016/j.etap.2016.05.014, 10.3390/pharmaceutics8010004]; 2) ‘inhibition by the same spectrum of compounds’ [10.1124/dmd.108.020552, 10.1016/j.toxlet.2016.09.004]; 3) ‘association between gene polymorphisms’ [10.1158/1535-7163.mct-10-0597, 10.21203/rs.3.rs-2650157/v1, 10.3389/fphar.2020.01241]; 4) ‘bioactivation of toxic metabolites’ [10.22438/jeb/41/3/MRN-1276, 10.1007/s00204-017-1935-8, 10.1007/s00204-024-03710-9, 10.1002/em.21912]; 5) ‘sequential phase I–II metabolism’ [10.1074/jbc.m808321200, 10.1111/bcpt.14013, 10.5246/jcps.2022.08.051]. From Table S1 it can be seen the presence of conditionally direct (‘sequential phase I–II metabolism’ and ‘bioactivation of toxic metabolites’) and to a lesser extent indirect (e.g. ‘association between gene polymorphisms’) functional associations between SULTs and CYPs that may support the hypothesis of PPIs between different representatives of the two multigene families. Bioinformatic analysis of proteomics and transcriptomics data showed that SULT-CYP PPI aspect has not yet been studied in detail (Figure S3, Supplementary material) and some SULT- and CYP-encoded genes demonstrated tissue co-expression patterns (Figure S4, Supplementary material). To summarize, we have not found any experimental confirmation or refutation of the hypothesis of protein-protein interactions between SULT1E1/2A1 and CYP3A4/3A5. The methodology for literature search and bioinformatic analysis of SULTs and CYP3A interaction data [10.1093/nar/gkaa1074, 10.1093/nar/gkz430, 10.1006/bbrc.1999.0960, 10.1093/nar/gkaa1074, 10.1007/s11192-009-0146-3, Dubaisi ST. Characterization of cytosolic sulfotransferase expression and regulation in human liver and intestine. Wayne State University Dissertations. 2019;2158. https://digitalcommons.wayne.edu/oa_dissertations/2158, 10.1124/dmd.119.086363, 10.1124/dmd.105.008672] is described in detail in the Supplementary Material.

Current literature and bioinformatic evidence does not support the existence of direct physical (protein–protein) interactions or stable complexes between the SULTs (SULT1E1 and SULT2A1) and the CYP3A enzymes (CYP3A4 and CYP3A5), as no studies have specifically addressed this question. Instead, available data point to their participation in sequential or parallel metabolic pathways acting on shared substrates, without confirmed direct molecular contact.

Thus, characterizing SULTs (SULT1E1 and SULT2A1) and CYP3A (CYP3A4 and CYP3A5) interactions could uncover new regulatory mechanisms, explain metabolic balance, and reveal mutual functional modulation between these enzymes. Such insights would be essential for understanding steroid homeostasis, drug metabolism, and interindividual variability in pharmacological responses associated with dysregulation of these enzymes in humans.

## 2. Materials and Methods

### 2.1. Instruments

For the quality control of protein purification MALDI-TOF Microflex LRF (Bruker Daltonik GmbH, Bremen, Germany) was used. Mass spectra acquisition and processing were performed using FlexControl 3.4 and FlexAnalysis 3.0 software (Bruker Daltonics GmbH, Germany), respectively.

The HPLC analysis of enzymatic reaction products was performed on an Agilent Technologies 1200 Series chromatograph equipped with an Agilent 6410 Triple Quadrupole LC/MS system (Agilent, Santa Clara, CA, USA).

PPI analysis was performed using SPR biosensor Biacore 3000 (Cytiva, Marlborough, MA, USA).

### 2.2. Protein Preparations

Highly purified (>95% according to SDS-PAGE) preparations of recombinant proteins CYP3A5, CYP3A4: the expression vectors CYP3A4_pCWori+HT and CYP3A5_pCWori+HT (with deleted hydrophobic N-terminal sequences—Δ3-12 for CYP3A4 and Δ2-12 for CYP3A5, a Ser-Thr-linker and 6xHis-tag added to the C-terminal sequence to facilitate purification); SULT1E1, and SULT2A1:the expression vectors SULT1E1 pCW-LIC, full-size, 6xHis-tag added to the N-terminal sequence to facilitate purification, SULT2A1 pCW-LIC, full-size, 6xHis-tag added to the C-terminal sequence to facilitate purification (Figure 1) were carried out in the Institute of Bioorganic Chemistry of the National Academy of Sciences of Belarus using molecular cloning and heterologous expression in a bacterial system (E. coli) followed by metal affinity and ion exchange chromatography for protein purification. These modifications and the purification procedure yield approximately 200 nmol of CYP3A4, 350 nmol of CYP3A5, 2000 nmol of SULT1E1, 1000 nmol of SULT2A1 per liter of expression medium with a purity of over 95% as determined by SDS-PAGE. The expression and purification protocols of CYP3A5, CYP3A4, SULT1E1, and SULT2A1 have been presented in our previous works.^14, 16^ The primary structures of the studied proteins are shown in Figure 2. MALDI-TOF spectra and SDS-PAGE images for the purity control of the proteins are presented in Figure 1.

**Fig. 1.**
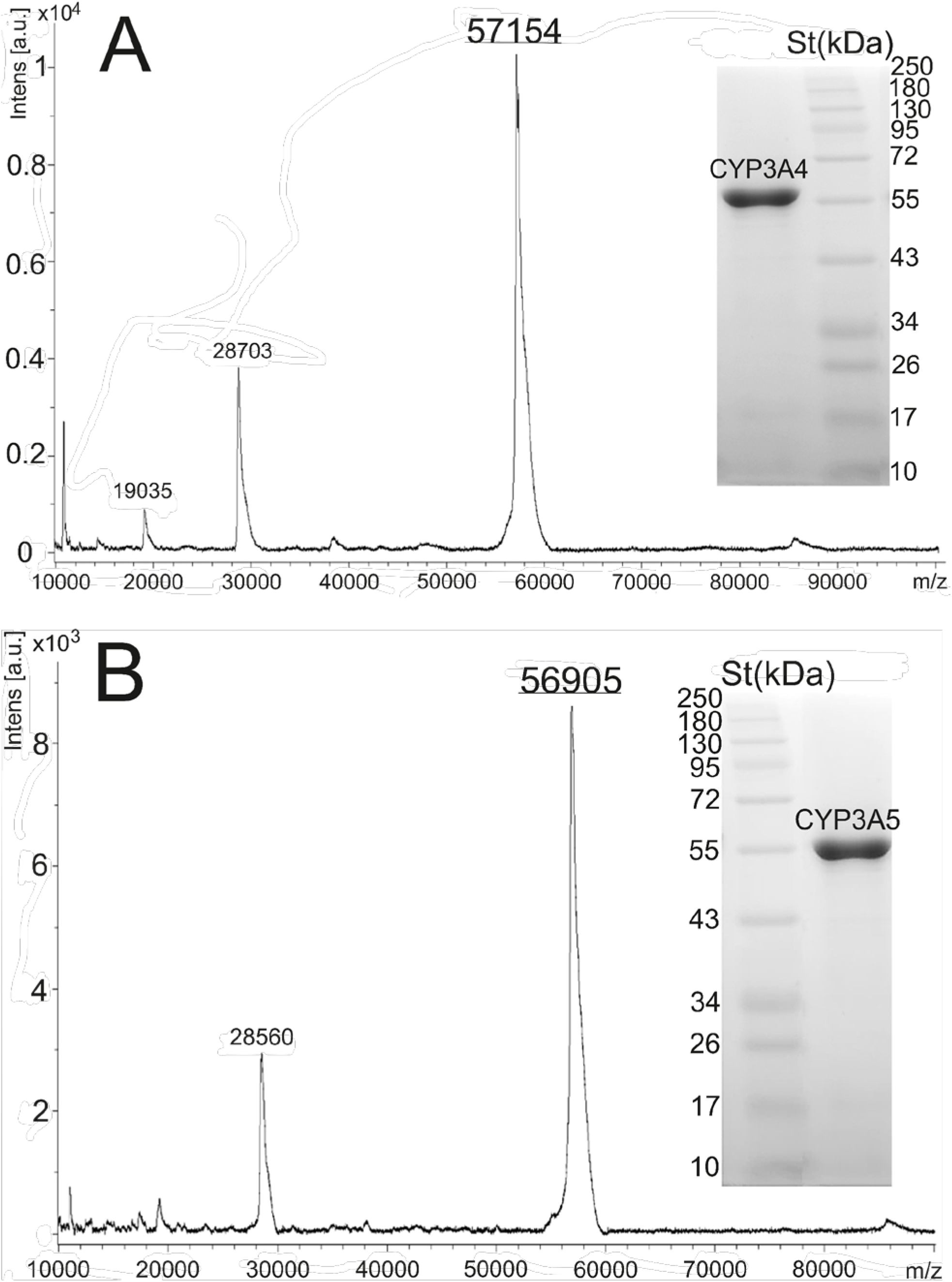
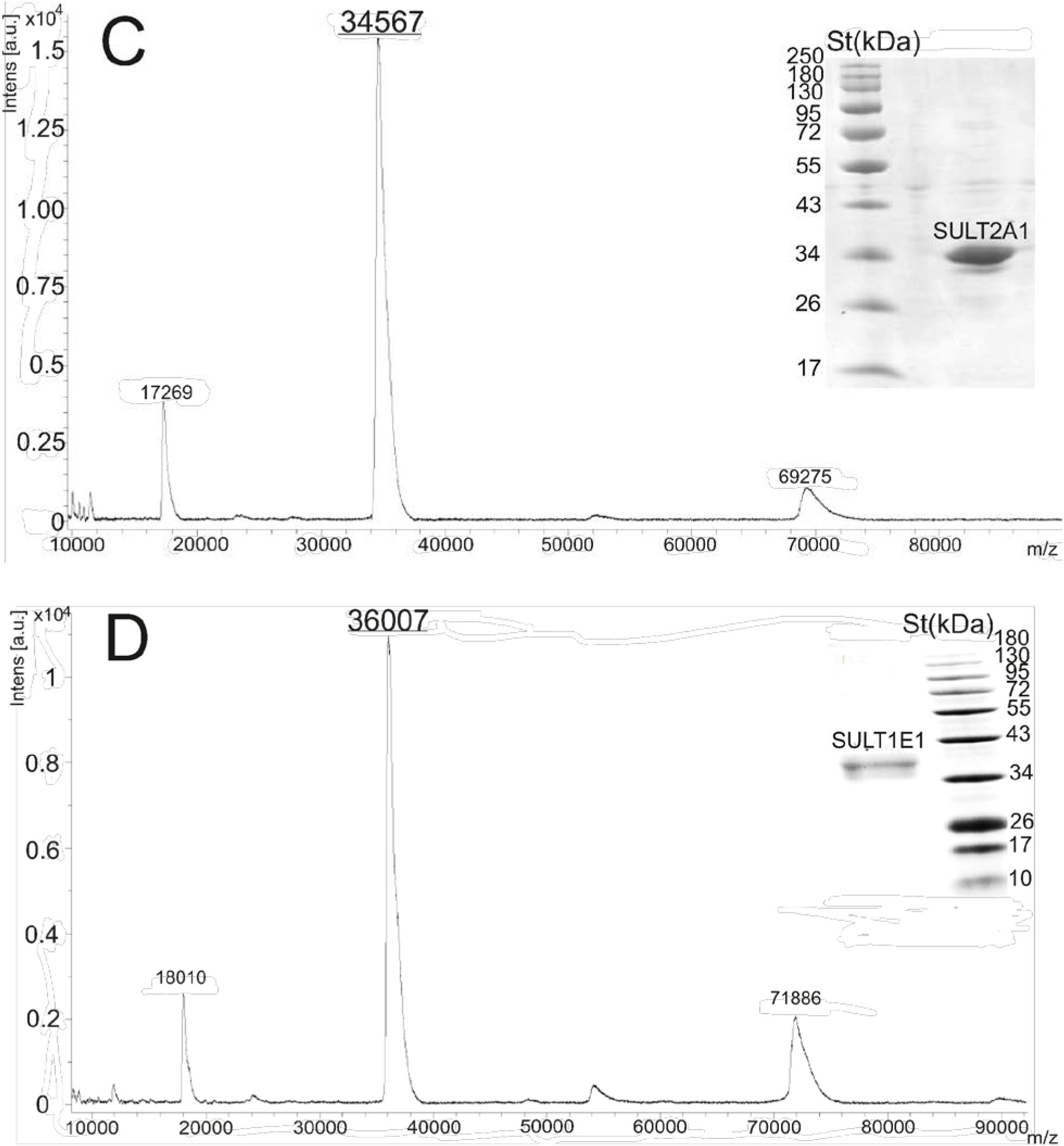
Protein purification data. A. Preparation of CYP3A4 with a mass of 57154 Da (according to MALDI-TOF mass spectrometry). SDS-PAGE analysis of CYP3A4. St. - protein molecular weight standard (250-10 kDa, Protein marker NEB #P7719S). B. Preparation of CYP3A5 with a mass of 56905 Da (according to MALDI-TOF mass spectrometry). SDS-PAGE analysis of CYP3A5. St. - protein molecular weight standard (250-10 Da, Protein marker NEB #P7719S). C. Preparation of SULT2A1 with a mass of 34567 Da (according to MALDI-TOF mass spectrometry). SDS-PAGE analysis of SULT2A1 protein. St. - protein molecular weight standard (250-10kDa, NEB #P7719S). D. Preparation of SULT1E1 with a mass of 36007 Da (according to MALDI-TOF mass spectrometry). SDS-PAGE analysis of SULT1E1 protein. St. - protein molecular weight standard (250-10kDa, NEB #P7719S).

**Fig. 2.**
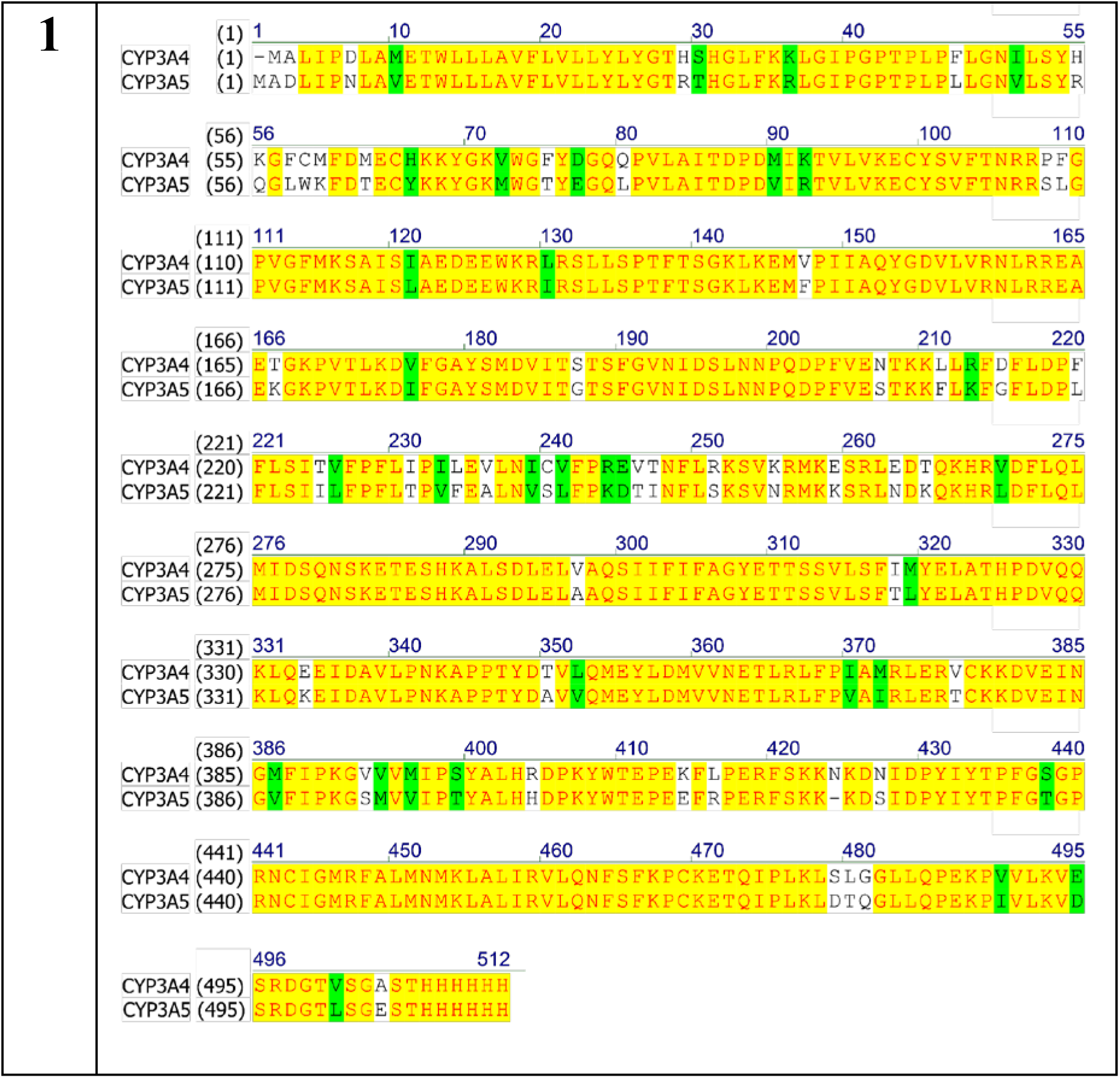
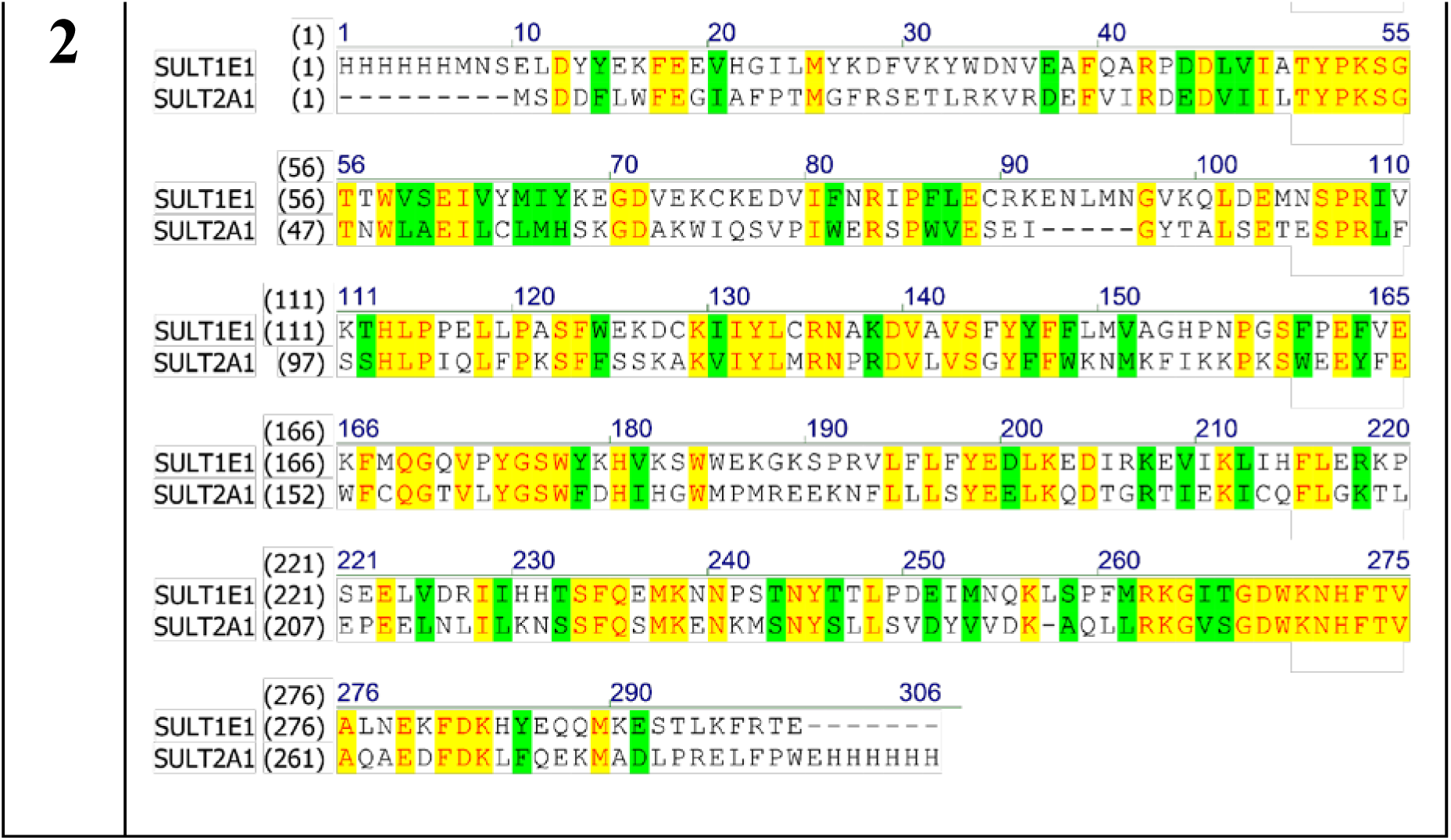
Comparison of amino acid sequences of the studied proteins using the Align software package (Invitrogen, USA). 1. CYP3A4 and CYP3A5; 2. SULT1E1 and SULT2A1. Green - the homologous part of the amino acid sequence; yellow - the identical part of the amino acid sequence.

### 2.3. Reagents

Dehydroepiandrosterone (DHEA), testosterone, NADPH and 3′-phosphoadenosine-5′- phosphosulfate (PAPS) were obtained from Sigma-Aldrich (St. Louis, MI, USA). HBS-EP+ buffer (10 mM HEPES, 150 mM NaCl, 3 mM EDTA, 0.05% P20 detergent, pH 7.4), 10mM sodium acetate buffers (pH 5.0 and 5.5), a reagent kit for the covalent immobilization of proteins via primary amino groups (1-ethyl-3-(3-dimethylaminopropyl)-carbodiimide-HCl (EDC) and N-hydroxysuccinimide (NHS)) were obtained from Cytiva (Marlborough, MA, USA). The remaining reagents were obtained from local suppliers.

### 2.4. Bioinformatic Analysis

#### 2.4.1. Protein–Protein Interaction Network Construction

Construction of PPI networks was performed using the STRING database v.11.5 (https://string-db.org, accessed 25 June 2026) with the following settings: meaning of network edges: evidence; active interaction sources: text mining, databases, co-expression, neighborhood, gene fusion, and co-occurrence; minimum required interaction score: highest confidence 0.4, max number of interactors to show: query proteins only; k-means clustering.^18^

#### 2.4.2. Co-Expression Analysis

The co-expression of genes encoding SULTs and CYPs (*SULT1E1-CYP3A4*, *SULT1E1-CYP3A5, SULT2A1-CYP3A4*, and *SULT2A1-CYP3A5*) in normal human tissues was analyzed using the GEPIA2 portal (http://gepia2.cancer-pku.cn/#correlation, accessed 08.21.2025).^19^ For this analysis, we selected tissues known to have high expression levels of target genes, namely: small intestine (terminal ileum), liver, adrenal gland, uterus, and mammary gland.^16,17,18^ The following cut-off criteria were applied for the analysis: a Spearman correlation coefficient values |R|>0.5, a number of clinical cases >30, and p-value <0.0001.

#### 2.4.3. Bibliometric analysis

The search for articles on non-steroidogenic CYPs and cytosolic sulfotransferases of the SULT family was performed in the PubMed and Scilit databases using the query “SULT and CYP”. The search depth was more than 20 years (2004–2025); article language – English; articles were sorted by the best match (relevance) and the content of each abstract was examined manually. Articles that mention any functional associations between SULTs and CYPs metabolizing exogenous or endogenous compounds were included in the final list.

Analysis of term co-occurrences and network visualization were performed using VOSviewer 1.6.20 with following settings: counting method – binary method; minimum number of occurrences of a term – 10. Of the 2401 initial terms, 38 met the threshold, terms with an uncertain meaning such as ‘study’, ‘patient’, ‘data’, ‘control’, ‘woman’, ‘role’, ‘year’, ‘combination’, ‘addition’, ‘confidence interval’, and ‘gene’ as well as duplicate terms (‘SULT1A1 gene’, ‘metabolite’, ‘sulfotransferase’, and ‘cytochrome’) were omitted. 26 terms were selected for further analysis.^20^

#### 2.4.4. Structural Prediction of Protein–Protein Complexes In Silico

Prediction of protein-protein complex structures with unknown spatial configurations was conducted using AlphaFold2 implemented on the ColabFold server v1.5.2 under default parameters. Canonical amino acid sequences were retrieved from the UniProt database.^21,22^ For each complex, five structural hypotheses were generated. Selection of representative models was based on the following thresholds: pLDDT (predicted local distance difference test) >70 and pTMscore (predicted template modeling score) ≥0.5.^23,24^ While pLDDT reflects the local folding reliability of the polypeptide chain, pTMscore estimates the quality of the protein-protein interface. Notably, some models with visually implausible geometries still satisfied these thresholds. The obtained models were post-processed using UCSF Chimera v1.16^28^, the heme was parametrized and transferred from PDB ID PDB ID 6MJM (CYP3A5) and 5VCC (CYP3A4) prior to energy minimization. To enhance selection rigor, the PDBePISA online tool^25^ was employed to evaluate interfacial characteristics, including contact area (>300 Å²) and the number of salt bridges and hydrogen bonds, with preference given to models exhibiting the most extensive interactions.^26,27^ Final models meeting all criteria were retained for analysis. Additionally, PDBePISA was used to identify specific amino acid residues contributing to the protein-protein interface, particularly those involved in salt bridge and hydrogen bond formation. Structural visualizations were generated using UCSF Chimera v1.16^28^.

Tunnel prediction within the SULT-CYP models was carried out with Caver Analyst v.2.0 using a probe diameter of 0.9 Å.^29^

### 2.5. Surface Plasmon Resonance Analysis

Real-time measurements of PPIs were performed at 25°C using CM5 optical chips coated with carboxymethyl-dextran (CM-dextran). PPI sensorgrams were recorded as the change in the biosensor signal in resonance units (RU) per unit of time (s). SULT2A1 and SULT1E1 were covalently immobilized onto the CM5 chip via amino groups. The carboxyl groups of CM-dextran were activated by a 1:1 (v/v) mixture of 0.4 M EDC and 0.1 M NHS for 7 min at a flow rate of 5 µL/min. Proteins (15 μg/mL) in 10 mM sodium acetate buffer (pH 5.0 for SULT2A1, pH 5.5 for SULT1E1) were injected for 2 min at a flow rate of 5 μL/min. Next, the chip surface was washed with HBS-EP+ buffer for 60 min.

CYP3A4 and CYP3A5 were injected as analytes in HBS-EP+ buffer at concentration ranges of 10-150 nM and 2.5-25 nM, respectively, through the control (intact CM-dextran) and working (with immobilized protein) channels for 2 min at 15 µL/min. The dissociation of protein-protein complexes was recorded at the same flow rate for 5 min. In addition, the same experiments were carried out in the presence of 3′-phosphoadenosine-5′-phosphosulfate – PAPS (up to 1 mM) to assess the PPI-modulating effect. The resulting set of sensorgrams was obtained by subtracting the biosensor signal in the control channel from the signal in the working channel and analyzed according to the “Heterogeneous ligand (parallel reactions)” model using the Biacore BIAevaluation software v. 4.1.1 (Cytiva, Marlborough, MA, USA).

The “Heterogeneous ligand (parallel reactions)” model describes each sensorgram as the sum of independent interaction reactions of a single analyte (CYP) with two types of ligands (S1 and S2):

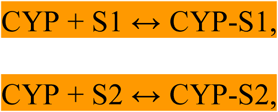

where k_on1_ and k_on2_ are the association rate constants, k_off1_ and k_off2_ are the dissociation rate constants.The resulting affinities (K_a1_ and K_a2_) are obtained as K_a1_= k_on1_/k_off1_ and K_a2_ = k_on2_/ k_off2_, respectively 30.

### 2.6. Biochemical Tests

#### 2.6.1. CYP3A4 and 3A5 Enzymatic Activity in the Presence of SULTs

The enzymatic activity of CYP3A4 and CYP3A5 was assessed in a reconstituted system containing 1 µM CYP3A4 or CYP3A5, 2 µM cytochrome P450 reductase of rat (CPR), the full-length rat enzyme, 2 µM microsomal cytochrome b5 (CYB5A), the full-length human protein with a C-terminal His-tag, 0.02 % sodium cholate, 20 µg of lipid mix (equal amounts of DOPC (1,2-Dioleoyl-sn-glycero-3-phosphocholine), DLPC (1,2-Dilauroyl-sn-glycero-3-phosphocholine), and PS(phosphatidylserine)) and 25 µM testosterone in a buffer comprising 50 mM HEPES (pH 7.4) and 10 mM MgCl_2_. To estimate the effect of SULTs on CYP3A4 and CYP3A5 activity, 1 µM SULT1E1 or SULT2A1 was added to the system. The molar ratio of CYP:SULT was 1:1.

After incubating the protein aliquots at 37 ^◦^C for 10 min, the reaction was started by adding by adding 250 µM NADPH. Then, the mixture was incubated at 37 ^◦^C for 30 min. The reaction was stopped by adding 5 mL of methylene chloride and subsequent centrifugation at 3000 rpm for 5 min. The upper organic layer was taken and evaporated in an argon atmosphere. After that, 100 µL of methanol was added to dissolve the precipitate.

Product formation was detected with HPLC using C18 Luna column (100 Å, 250 *×* 4.6 mm) and Agilent 1260 Infinity III (Agilent Technologies, Germany). Column elution was performed at a flow rate of 1 mL/min using a linear gradient from mobile phase A (62 % methanol) to mobile phase B (100 % methanol). The eluate was analyzed at a wavelength of 254 nm. The column temperature was 37 °C.

#### 2.6.2. SULTs Enzymatic Activity in the Presence of CYP3A4 and 3A5

The enzymatic activity of SULTs was assessed in a reconstituted system containing 2 µM SULT and 60 µM PAPS in a 50 mM Tris-HCl buffer (pH 7.5) comprising 10 mM MgCl_2_. To test the effect of CYP3A4 and CYP3A5 on the catalytic activity of the studied SULTs, 2 µM CYP3A4 or CYP3A5 was added to the system. The molar ratio of CYP:SULT was 1:1. After incubating the protein aliquots at 37 *^◦^*C for 10 min, the reaction was started by adding 50 µM DHEA with subsequent incubation at 37 *^◦^*C for 3 hours. The reaction was stopped by adding 5 mL of methylene chloride and subsequent centrifugation at 3000 rpm for 5 min. The upper organic layer was taken and evaporated in an argon atmosphere. After that, 100 µL of acetonitrile was added to dissolve the precipitate.

The products of the enzymatic reaction of SULTs, DHEA-S, was analyzed with LC-MS using Zorbax Eclipse XDB-C18 column (80 Å, 150 x 4.6 mm, 5µm) and Agilent Technologies 1200 LC – 6410B Triple Quadrupole system (Santa Clara, CA, USA). Mobile phase A was 0.1 vol % formic acid in water and mobile phase B was 0.1 vol % formic acid in mixture isopropanol: methanol (1:3). Isocratic elution 30 % phase B during 1 min, linear gradient 30–100 % phase B in 1–10 min. The flow rate was 0.5 mL×min^-1^, and the column temperature was 40 ^◦^C. Injection volume – 20 µL. Stop time – 20 min. ASL temperature +10°С Chromatographic separation was followed by APCI ionization in positive ion detection mode with next parameters: gas temperature + 200 ^◦^C; gas flow drying gas – 8 L/min; nebulizer – 40 psi; capillary voltage – 4000 V; corona current - 4µA; vaporizer + 400 ^◦^C; fragmentor voltage – 80 V; mass range – 100-1500 m/z in the Scan mode.

For quantitative assay extracted ion chromatograms (EIC) for ions with m/z 271.2 for DHEA-S and DHEA were used. Solutions of DHEA-S and DHEA with concentrations of 100 and 200 µM in methanol were used as external standards.

## 3. Results

### 3.1. Systems Biology Analysis of SULT–CYP3A Associations

#### 3.1.1. Bibliometric analysis

A visualization of MeSH keyword co-occurrence in articles is shown in Figure 3 (a map of term co-occurrence in the article’s abstracts). At least two main clusters (blue and purple) of terms can be distinguished. The first cluster reflects studies devoted to identifying associations of SULT and CYP gene polymorphisms with the disease prognosis, drug treatment efficiency, and ageing. The terms ‘SULT1A1’ and ‘SULT1A1*2’ genes (the most common allelic variant of SULT1A1 in the human population) significantly prevail over other SULTs in the texts analysed. The second cluster includes terms reflecting studies devoted to compound metabolism from the point of view of the functional activity of cytosolic SULTs and microsomal CYPs with broad substrate specificity (CYP2D6, CYP3A4, and CYP3A5). It is possible to distinguish the co-occurrence of ‘SULT1A1 and CYP1B1’, ‘SULT1A1 and CYP1A1’ (the first cluster); ‘SULT1A1 and CYP2D6’, ‘SULT1A1 and CYP3A5’, ‘SULT1A1 and CYP3A4’ (the second cluster). Interestingly, co-occurrence of the terms ‘SULT1A1’, ‘CYP1A1’ is shown in the context of studies on gene expression profiling. A more detailed map of co-occurrence of different representatives of non-steroidogenic CYPs and cytosolic sulfotransferases is shown in Figure 4, from which it is evident that there are associations between 12 CYPs (CYP1A1, 1A2, 1B1, 2B6, 2D6, 2C9, 2C19, 2E1, 3A4, 3A5, 7A1, 8B1) and 5 SULTs (SULT1A1, 1A2, 1B1, 2A1, 2B1).

**Fig. 3.**
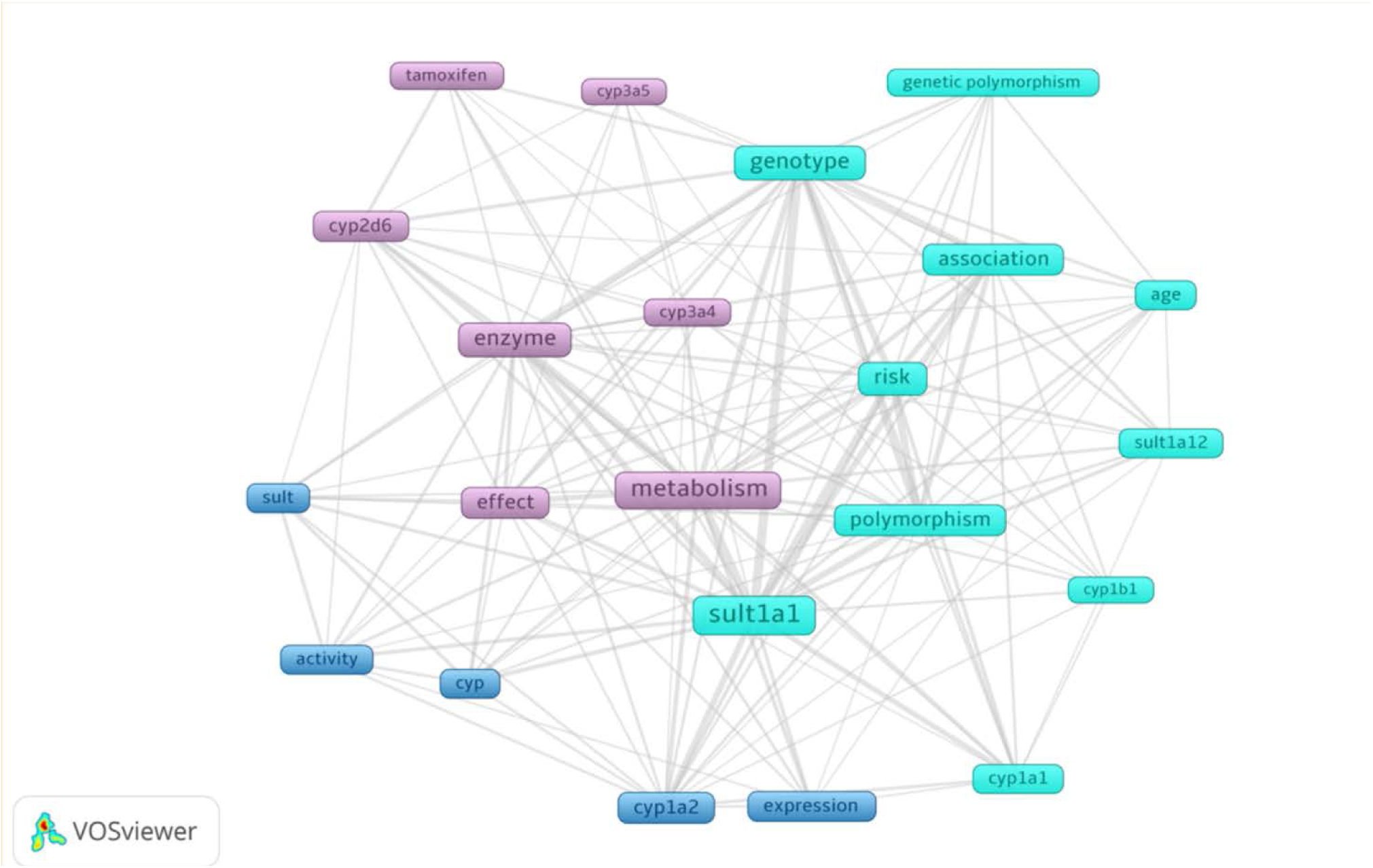
Co-occurrence map of specific biomedical terms in 77 articles abstracts. The analysis included terms with a minimum of 10 occurrences. Network visualization was performed using VOSviewer 1.6.20.

**Fig. 4.**
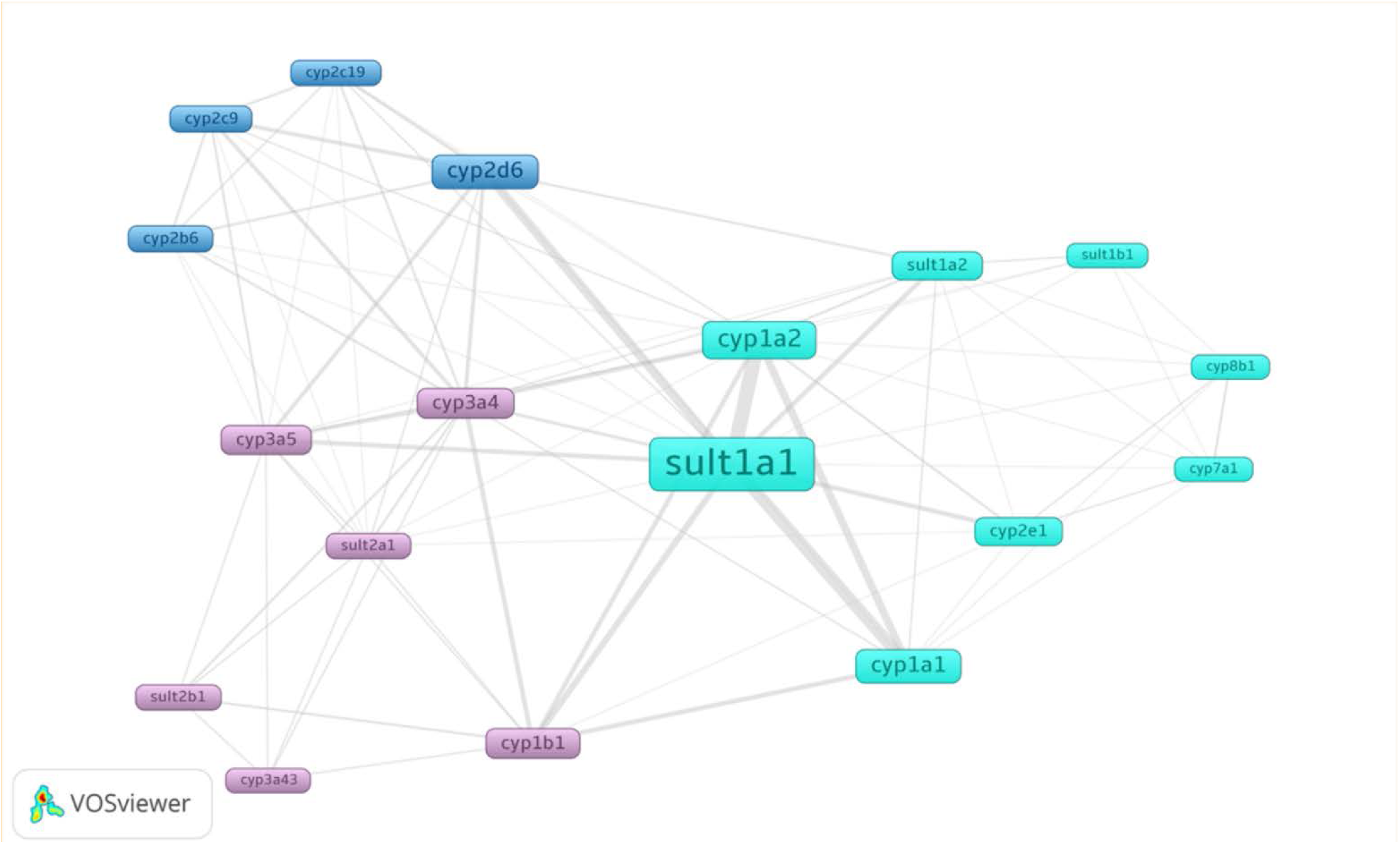
Co-occurrence map of SULTs and CYPs in 77 articles abstracts. The analysis included terms with a minimum of 10 occurrences. Network visualization was performed using VOSviewer 1.6.20.

#### 3.1.2. Literature data analysis

The analysis of the texts of the selected articles revealed 5 characteristic types of functional associations between SULTs and non-steroidogenic microsomal CYPs: 1) ‘gene co-expression under disease conditions’, 2) ‘inhibition by the same spectrum of compounds’, 3) ‘association between gene polymorphisms’, 4) ‘bioactivation of toxic metabolites’, 5) ‘sequential phase I–II metabolism’ (Table 1). From Table 1 it can be seen the presence of conditionally direct (‘sequential phase I–II metabolism’ and ‘bioactivation of toxic metabolites’) and to a lesser extent indirect (e.g. ‘association between gene polymorphisms’) functional associations between SULTs and non-steroidogenic microsomal CYPs. Functional associations between SULT and CYPs may support the hypothesis of PPIs between different representatives of the two multigene families. Of note, the PPI aspect has not yet been studied in detail, and we have not found any experimental confirmation or refutation of this hypothesis.

**Table 1.**
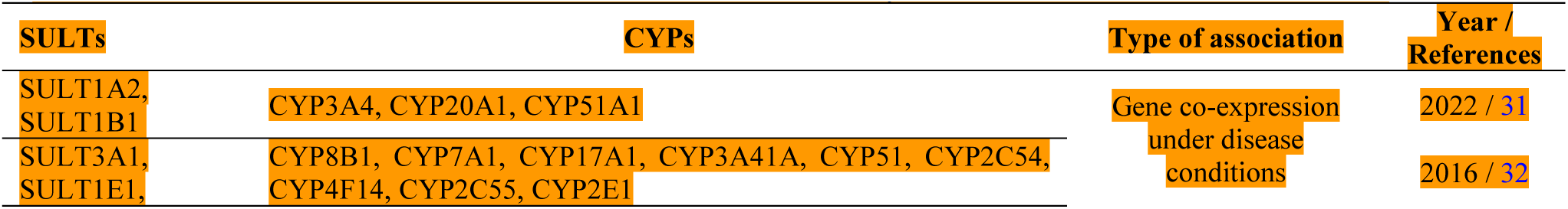
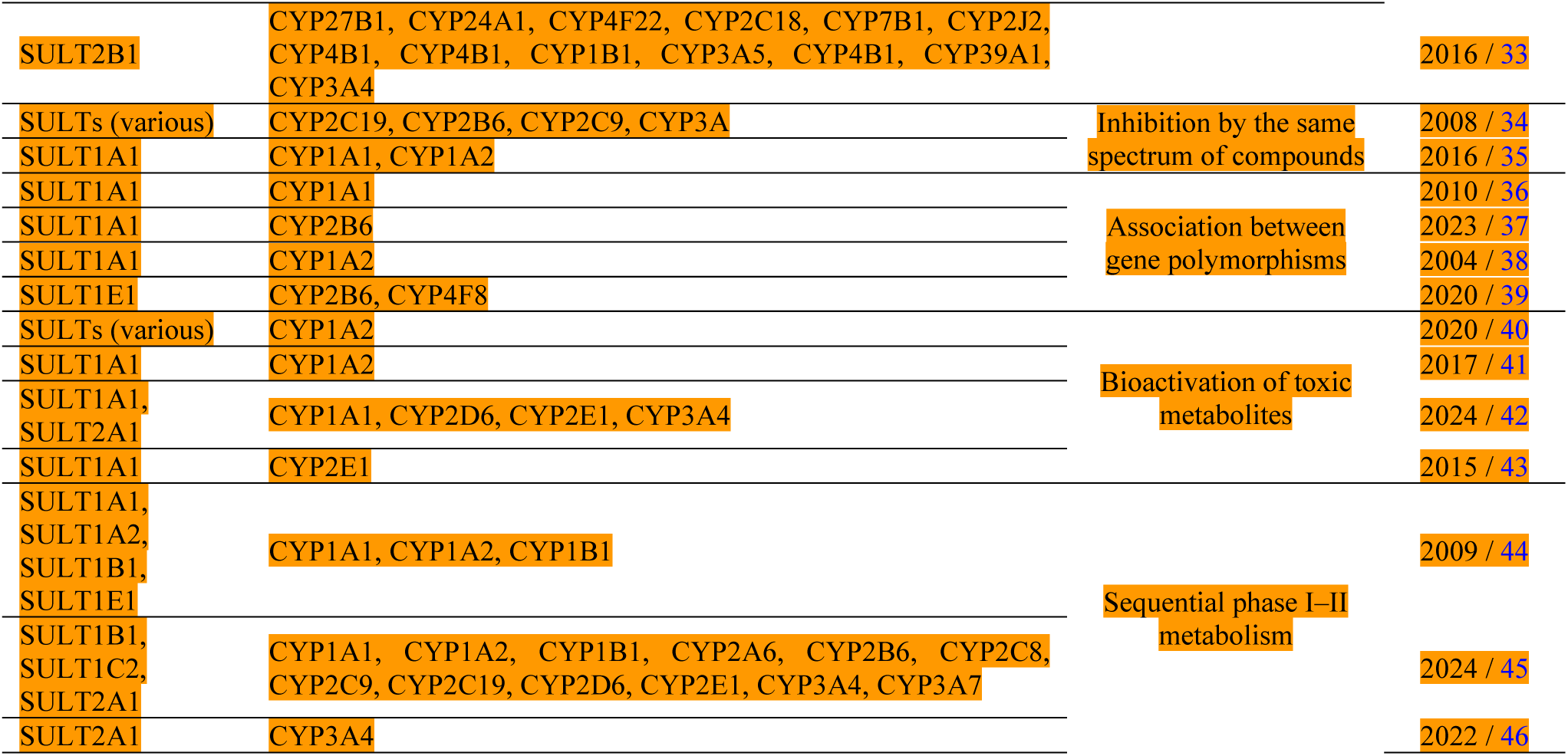
Functional associations between SULTs and CYPs enzymes based on the literature data.

#### 3.1.3. Protein-protein interactions

To investigate potential associations between SULT1E1/SULT2A1 and CYP3A4/CYP3A5, an analysis of possible protein–protein interactions (PPIs), gene co-expression patterns was conducted. Using STRINGdb data, a PPI network was generated for the target proteins (Figure 5). The protein–protein association network of our proteins, indicates their association based on terms co-occurrence in literature, curated database annotations (DIP (Database of Interacting Proteins), KEGG (Kyoto Encyclopedia of Genes and Genomes), GO (Gene Ontology), BioGRID, IntAct, MINT, PubMed, OMIM), and tissue-specific co-expression, thereby providing a rationale for subsequent in-depth analysis.

**Fig. 5.**
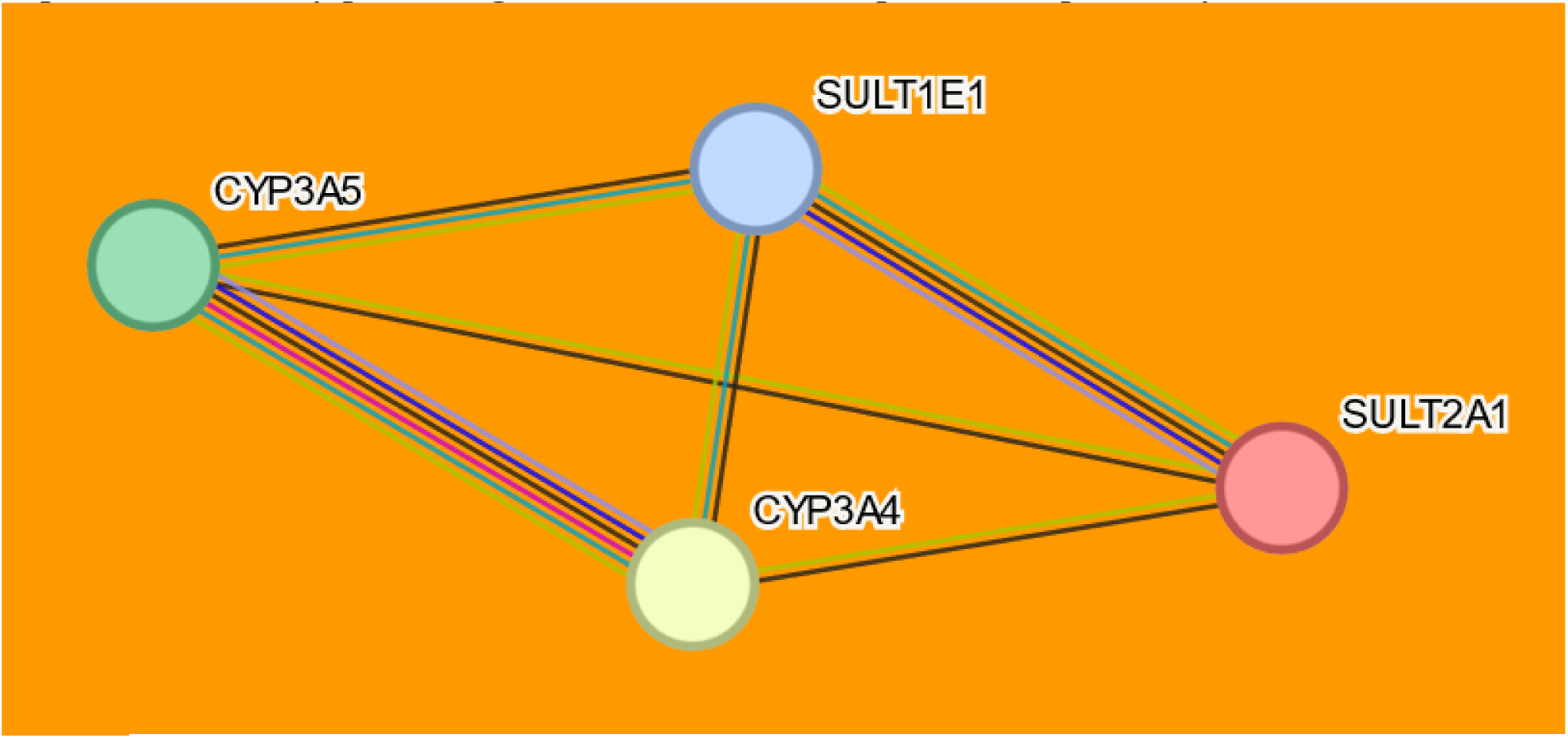
Protein–protein association network involving SULTs and CYPs according to the STRINGdb data. The edges between the nodes correspond to known PPI: from curated databases (cyan), experimentally determined (magenta); predicted PPI: gene neighborhood (green), gene co-occurrence (blue); other associations: co-expression (black), protein homology (purple).

#### 3.1.4. Co-expression analysis

Gene co-expression analysis of the SULTs of interest (SULT1E1 and SULT2A1) and the microsomal CYPs (CYP3A4 and CYP3A5) was performed in tissues with the highest known expression levels of these enzymes — the small intestine, liver, adrenal glands, uterus, and mammary gland.^47,48,49^ The analysis revealed a moderate correlation (R˂0.70) for SULT1E1– CYP3A4 and strong correlations (R˃0.70) for SULT1E1–CYP3A5, SULT2A1–CYP3A4,and SULT2A1–CYP3A5 in the small intestine. A moderate correlation was also observed for SULT1E1–CYP3A4 expression in the liver (Figure 6).

**Fig. 6.**
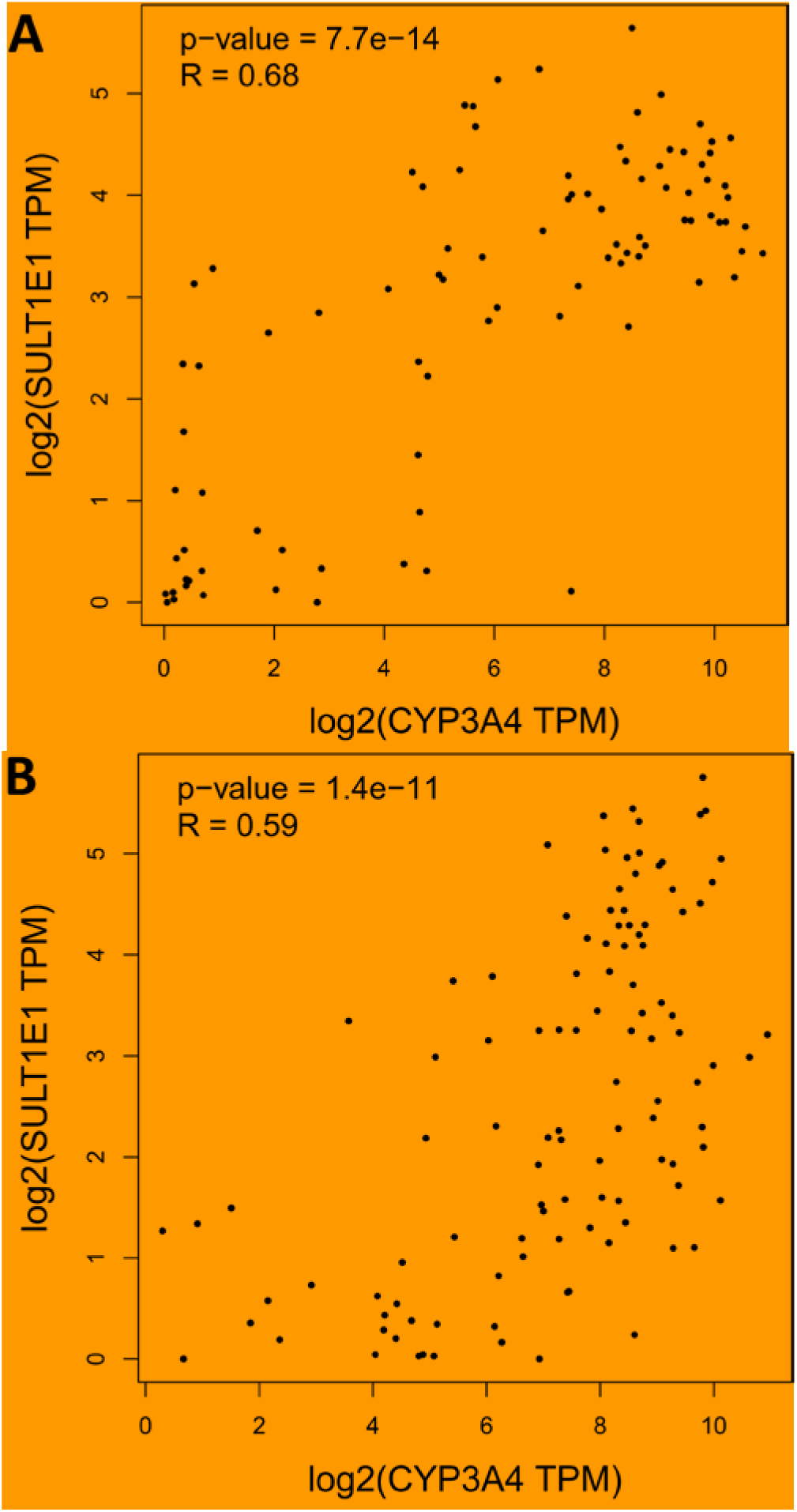
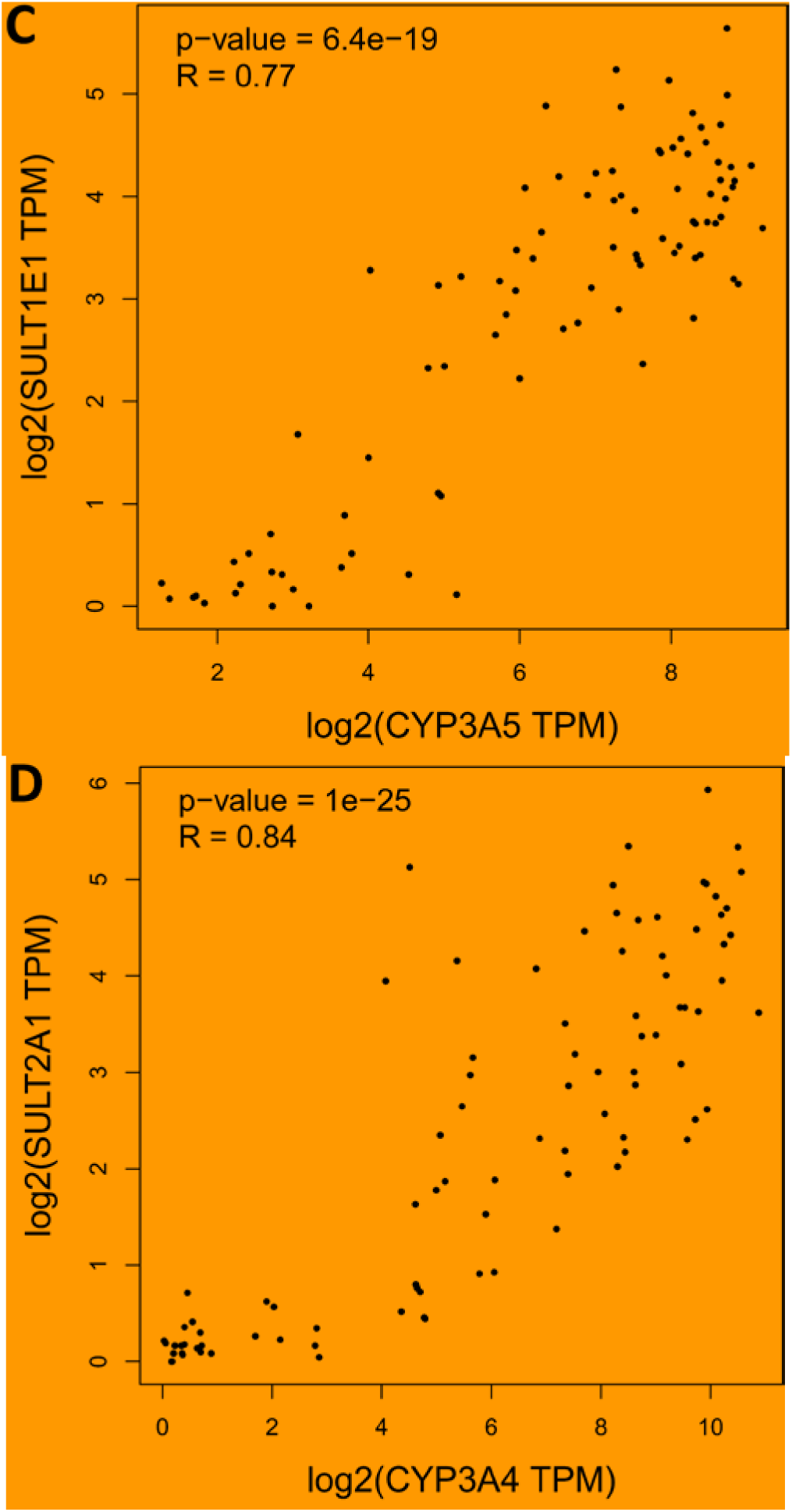
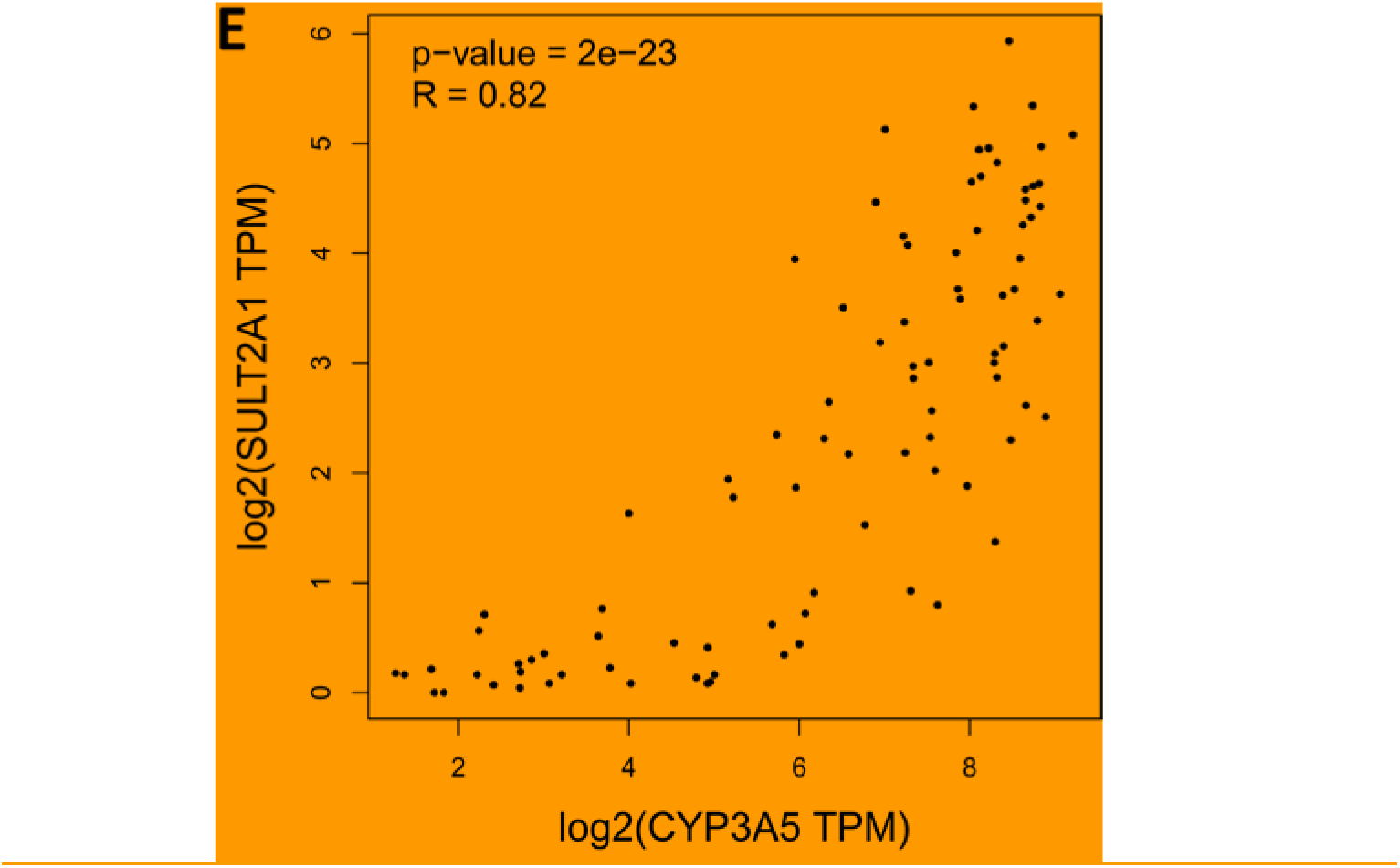
Gene co-expression for SULTs and CYP3A pairs. Scatter plots show the correlation of *SULT1E1–CYP3A4* expression in the small intestine (A) and liver (B), *SULT1E1–CYP3A5* (C), *SULT2A1–CYP3A4* (D), and *SULT2A1–CYP3A4* (E) in the small intestine. Spearman correlation coefficients (R) are shown on each plot. The analysis was performed using the GEPIA2 web-based tool and the original output images are shown.

In summary, systems biology analysis of SULT–CYP3A associations showed that no evidence of direct complex formation or physical PPI between the SULTs (SULT1E1 and SULT2A1) and the CYP3A enzymes (CYP3A4 and CYP3A5) was identified. However, literature data, gene co-expression analysis, and bibliometric analysis suggest the potential for such PPIs may exist.

### 3.2. SPR analysis of CYPs and SULTs interactions

To test the hypothesis of direct interactions between the SULTs (SULT1E1 or SULT2A1) and the CYP3A enzymes (CYP3A4 or CYP3A5), surface plasmon resonance (SPR) analysis was performed. Figure 7 shows representative sensorgrams of the resulting binary PPIs, which were used to determine the kinetic parameters (k_on_, k_off_) and equilibrium dissociation constants (K_d_) presented in Table 2. The effect of PAPS (3′-phosphoadenosine-5′-phosphosulfate) on complex formation was also evaluated.

**Fig. 7.**
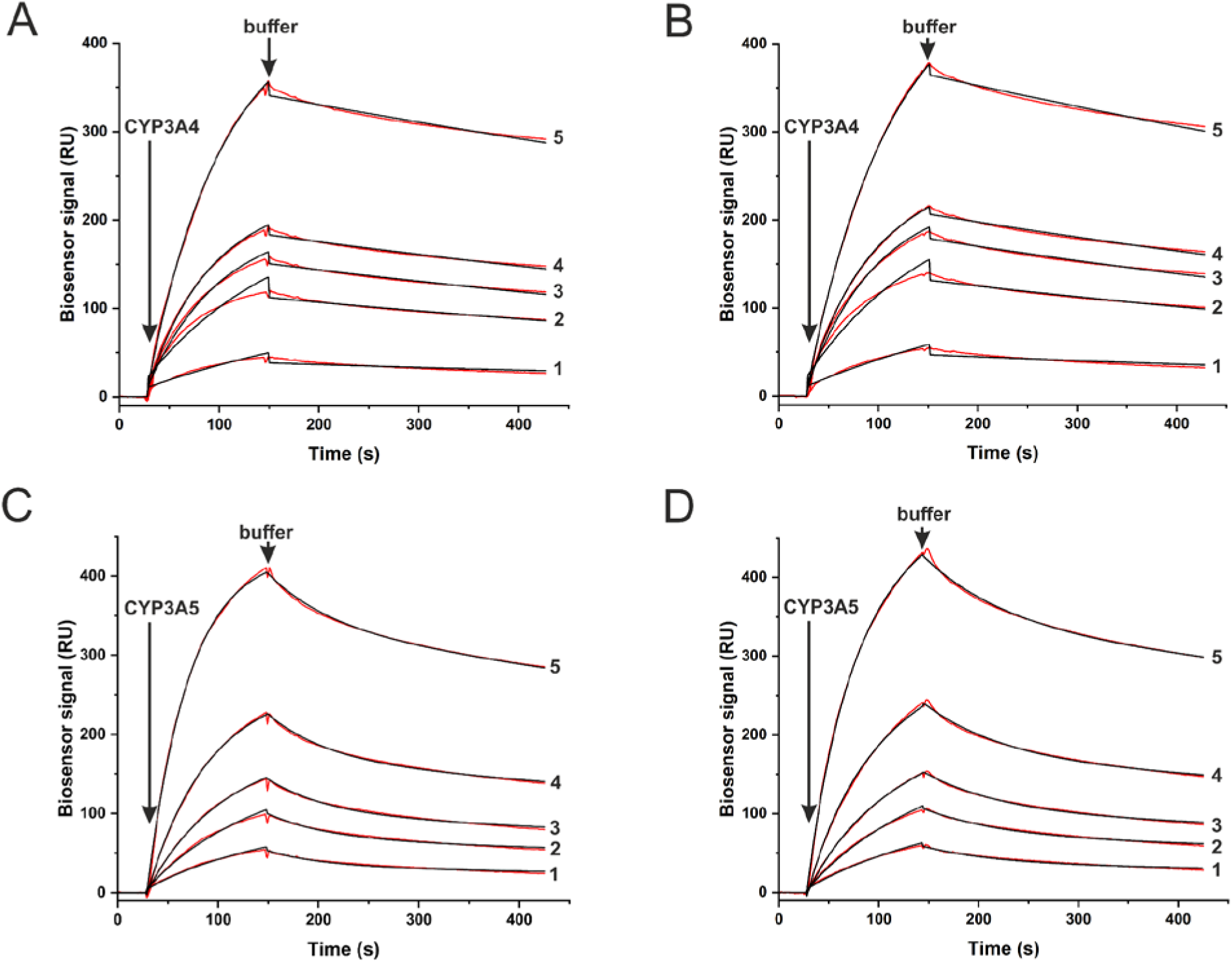
Sensorgrams depicting the binding of CYP3A4 (A, B) and CYP3A5 (C, D) to immobilized SULT2A1 (A, C) and SULT1E1 (B, D). Experimental data (red lines) were collected at concentrations of 10 (1), 25 (2), 50 (3), 75 (4), and 150 nM (5) of CYP3A4, and 2.5 (1), 5 (2), 10 (3), 15 (4), and 25 nM (5) of CYP3A5. The corresponding theoretical fitting curves are overlaid in black.

**Table 2.** Values of k_on_ (association rate constant), k_off_ (dissociation rate constant), and K_a_ (affinity) for the interactions of CYP3A4 and CYP3A5 with SULT1E1 and SULT2A1.

| Interaction Proteins (Ligand-Analyte) | $k_{on1}, M^{-1}s^{-1}$ | $k_{off1}, s^{-1}$ | $k_{on2}, M^{-1}s^{-1}$ | $k_{off2}, s^{-1}$ | $K_{a1}, M^{-1}$ | $K_{a2}, M^{-1}$ |
| --- | --- | --- | --- | --- | --- | --- |
| SULT2A1-CYP3A4 | $(8.44 \pm 0.02) \cdot 10^4$ | $(6.15 \pm 0.02) \cdot 10^{-4}$ | $(2.620 \pm 0.002) \cdot 10^5$ | $(9.59 \pm 0.03) \cdot 10^{-4}$ | $(1.37 \pm 0.01) \cdot 10^8$ | $(2.73 \pm 0.01) \cdot 10^8$ |
| SULT2A1-CYP3A4 (PAPS) | $(7.56 \pm 0.03) \cdot 10^4$ | $(6.35 \pm 0.04) \cdot 10^{-4}$ | $(1.990 \pm 0.004) \cdot 10^5$ | $(1.290 \pm 0.05) \cdot 10^{-3}$ | $(1.19 \pm 0.01) \cdot 10^8$ | $(1.54 \pm 0.01) \cdot 10^8$ |
| SULT2A1-CYP3A5 | $(8.95 \pm 0.16) \cdot 10^5$ | $(6.410 \pm 0.001) \cdot 10^{-4}$ | $(4.9 \pm 0.7) \cdot 10^5$ | $(1.65 \pm 0.02) \cdot 10^{-2}$ | $(1.4 \pm 0.03) \cdot 10^9$ | $(2.97 \pm 0.42) \cdot 10^7$ |
| SULT2A1-CYP3A5 (PAPS) | $(1.40 \pm 0.04) \cdot 10^6$ | $(2.43 \pm 0.04) \cdot 10^{-2}$ | $(5.92 \pm 0.06) \cdot 10^5$ | $(1.11 \pm 0.01) \cdot 10^{-3}$ | $(5.8 \pm 0.2) \cdot 10^7$ | $(5.33 \pm 0.07) \cdot 10^8$ |
| SULT1E1-CYP3A4 | $(2.810 \pm 0.003) \cdot 10^5$ | $(1.030 \pm 0.003) \cdot 10^{-3}$ | $(7.49 \pm 0.03) \cdot 10^4$ | $(6.95 \pm 0.03) \cdot 10^{-4}$ | $(2.72 \pm 0.01) \cdot 10^8$ | $(1.08 \pm 0.01) \cdot 10^8$ |
| SULT1E1-CYP3A4 (PAPS) | $(6.54 \pm 0.03) \cdot 10^4$ | $(6.50 \pm 0.03) \cdot 10^{-4}$ | $(2.04 \pm 0.004) \cdot 10^5$ | $(1.2 \pm 0.005) \cdot 10^{-3}$ | $(1.01 \pm 0.01) \cdot 10^8$ | $(1.69 \pm 0.01) \cdot 10^8$ |
| SULT1E1-CYP3A5 | $(1.26 \pm 0.03) \cdot 10^6$ | $(1.25 \pm 0.03) \cdot 10^{-2}$ | $(5.98 \pm 0.07) \cdot 10^5$ | $(5.1 \pm 0.2) \cdot 10^{-4}$ | $(1.01 \pm 0.03) \cdot 10^8$ | $(1.17 \pm 0.05) \cdot 10^9$ |
| SULT1E1-CYP3A5 (PAPS) | $(1.26 \pm 0.03) \cdot 10^6$ | $(1.62 \pm 0.03) \cdot 10^{-2}$ | $(1.63 \pm 0.02) \cdot 10^5$ | $(5.5 \pm 0.1) \cdot 10^{-4}$ | $(7.8 \pm 0.3) \cdot 10^7$ | $(2.96 \pm 0.06) \cdot 10^8$ |

PAPS final concentration is equal to 1 mM. Data are presented as the mean ± standard deviation (SD) of n=3 technical replicates.

### 3.3. Mutual Effects of CYPs and SULTs on Their Enzymatic Activities

To assess the functional relevance of the identified PPI, the mutual effects of the studied SULTs and CYP3A enzymes on each other’s catalytic activity were evaluated (Tables 3 and 4). It is worth noting that, in most cases, the presence of PAPS did not affect the activity of CYP3A4 or CYP3A5 under the tested conditions (Tables 3).

**Table 3.** Enzymatic activity of CYP3A enzymes in the presence of SULTs.

|  | **CYP3A4**<br>**Activity (min<sup>-1</sup>)** | **Resume** | **CYP3A5**<br>**Activity (min<sup>-1</sup>)** | **Resume** |
| --- | --- | --- | --- | --- |
| Control | 0.18±0.0001 | - | 0.06±0.0151 | - |
| SULT1E1 | 0.10±0.0152 | ↓* | 0.10±0.0095 | ↑ |
| SULT1E1/PAPS | 0.19±0.0214 | NC | 0.10±0.0033 | ↑ |
| SULT2A1 | 0.16±0.0632 | NC | 0.08±0.0091 | NC |
| SULT2A1/PAPS | 0.11±0.0094 | ↓ | 0.08±0.0188 | NC |

\*↑—slightly increased; ↓—slightly decreased; NC—not changed.

**Table 4.** Enzymatic activity of SULTs in the presence of CYP3A enzymes.

|  | **SULT1E1**<br>**Activity (min<sup>-1</sup>)** | **Resume** | **SULT2A1**<br>**Activity (min<sup>-1</sup>)** | **Resume** |
| --- | --- | --- | --- | --- |
| Control | 0.010±0.007 | - | 0.326±0.005 | - |
| CYP3A4 | 0.013±0.005 | NC | 0.451±0.014 | ↑* |
| CYP3A5 | 0.019±0.003 | NC | 0.679±0.022 | ↑↑ |

\*↑↑—increased; ↑—slightly increased; NC—not changed.

CYP3A4 activity decreased slightly in the presence of SULT or remained unchanged. In contrast, CYP3A5 activity slightly increased in reactions in the presence of SULTs.

When assessing SULT activity, the presence of CYP3A enzymes in the system contributed to a moderate increase in SULT2A1 activity and had no significant effect on SULT1E1 activity. The activity of SULT2A1 increased by 38% in the presence of CYP3A4, while the addition of CYP3A5 led to a 108% increase.

### 3.4. In Silico Modeling of Protein Complexes

Given that SPR analysis confirmed direct interactions between the CYP3A isoforms (CYP3A4 and CYP3A5) and the SULT isoforms (SULT2A1 and SULT1E1), we next sought to predict the structures of these protein complexes. We employed AlphaFold2 MMseqs2 for *in silico* modeling of the four possible complexes: CYP3A4–SULT2A1, CYP3A4–SULT1E1, CYP3A5–SULT2A1, and CYP3A5–SULT1E1. In addition to the standard AlphaFold2 confidence metrics, models were evaluated using interaction parameters derived from PDBePISA. Simulations were performed for both dimeric (‘dim’) and monomeric (‘mon’) forms of SULT enzymes, as the functional relevance of SULT oligomerization *in vivo* remains unclear.

No reliable models meeting quality thresholds or visual plausibility criteria could be generated for the CYP3A5–monSULT1E1, CYP3A5–dimSULT1E1, CYP3A4–monSULT2A1, or CYP3A4–dimSULT1E1 complexes; these were therefore excluded from further analysis. Ultimately, high-confidence structural models were obtained for four complexes (Table 5) involving CYP3A4 or CYP3A5 bound to the SULT enzymes of interest (Figure 8). The model were uploaded to the ModelArchive [10.1016/j.jmb.2025.168996] (DOIs: 10.5452/ma-1d4cm, 10.5452/ma-2f6fv, 10.5452/ma-i9o08, 10.5452/ma-i9o08). The analysis of protein–protein interfaces in the complexes revealed that the interacting residues of the SULT and CYP enzymes were located outside the heme- and substrate-binding sites, though often in close proximity to these regions. Comparative analysis of interfaces of dimeric SULT2A1 in its complexes with CYP3A4 and CYP3A5 showed that this sulfotransferase has common interaction sites with cytochromes of interest, concentrated in the middle and end of the SULT chain: here are observed such zones as [Lys-135, Asn-136, Lys-138, Phe-139, Ile-140, Lys-142, Lys-144], [Val-236, Asp-237] and [Val-240, Asp-241]. Lys-138 forms a salt bridge with both cytochromes of interest, and Lys-144 and Val-240 form hydrogen bonds with them. In the predicted models the described interacting regions of amino acids were located close to each other in the tertiary structure. And also, the CYP3A4– SULT1E1mon and CYP3A4–SULT2A1mon complexes are available for comparison of interfaces from the cytochrome (CYP3A4) side. According to the predicted data, CYP3A4 has the following identical interaction sites with both monomeric sulfotransferases of interest: Val-95, [Ser-131, Leu-132, Pro-135, Thr-138], [Ser-281, Glu-283, Thr-284, Glu-285], Tyr-430, [Gly-438, Pro-439], Arg-446. There are no salt bridges or hydrogen bonds among them.

**Fig. 8.**
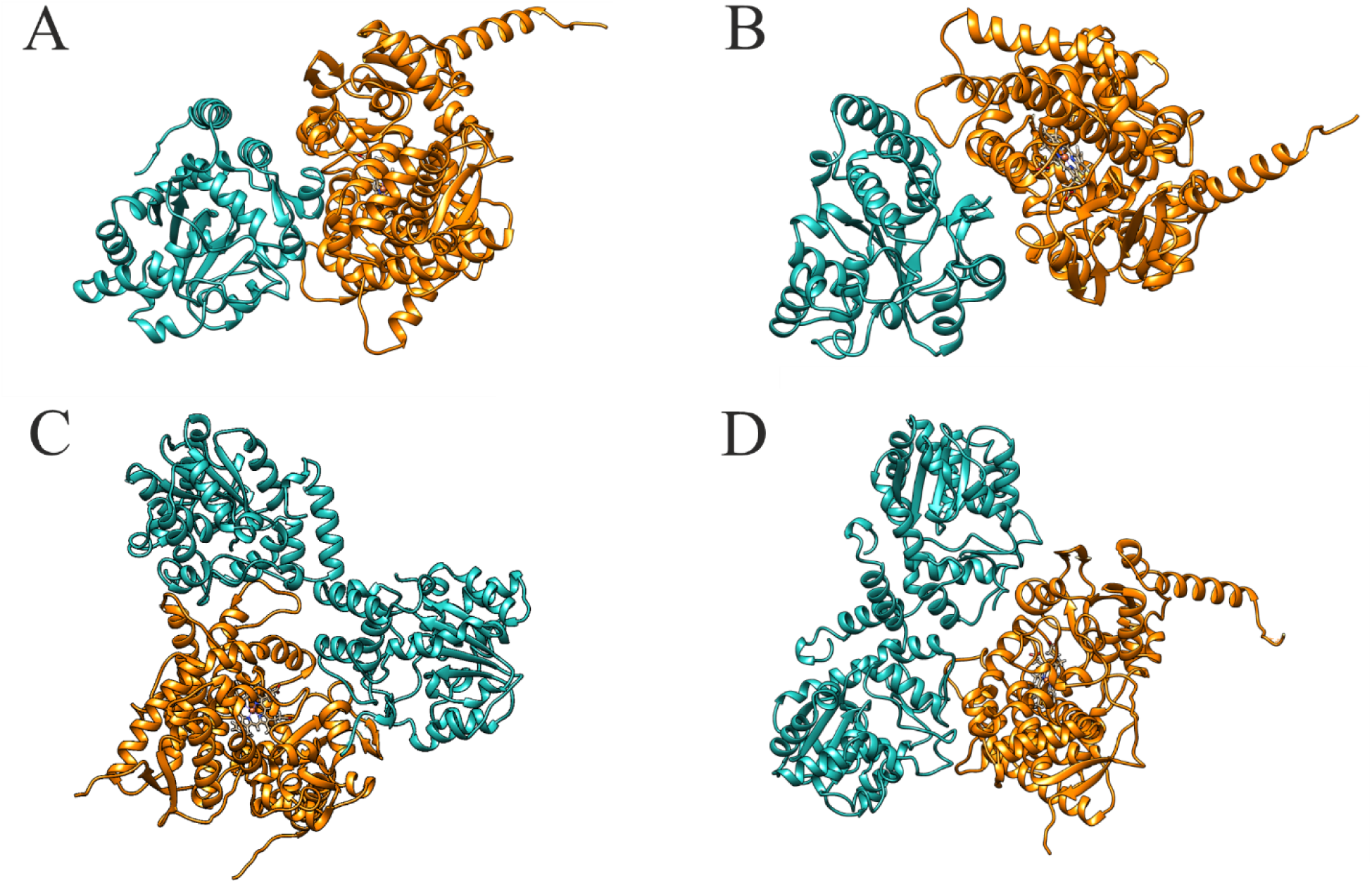
Models of the CYP3A4–monSULT1E1 (A), CYP3A4–monSULT2A1 (B), CYP3A4–dimSULT2A1 (C), and CYP3A5–dimSULT2A1 (D) complexes predicted in the AlphaFold 2 software. SULTs are shown in cyan and CYPs in orange.

**Table 5.** AlphaFold2 quality criteria and the structural characteristics of the intermolecular interface of protein–protein complexes of SULT2A1 and SULT1E1 with CYP3A4 and CYP3A5.

| Complexes | AlphaFold2 | PDBePISA |
| --- | --- | --- |

|  | pLDDT* | pTM** | Interface<br>area (Å<sup>2</sup>) | ΔG, kJ | H-Bonds<br>(HB) | Salt bridges<br>(SB) |
| --- | --- | --- | --- | --- | --- | --- |
| CYP3A4-mon***SULT2A1 | 86 | 0.7 | 715 | -1.3 | 5 | 5 |
| CYP3A4-dim****SULT2A1 | 86 | 0.5 | 1856 | -9.1 | 15 | 3 |
| CYP3A4-monSULT1E1 | 84 | 0.7 | 1242 | -4.4 | 7 | 9 |
| CYP3A5-dimSULT2A1 | 84 | 0.5 | 1092 | -4.3 | 6 | 4 |

\*-pLDDT (predicted local distance difference test), \*\*-pTMscore (predicted template modeling score), \*\*\*-mon (monomer), \*\*\*\*-dim (dimer).

In addition, we used Caver Analyst v.2.0 to predict potential substrate access tunnels in the modeled CYP–SULT complexes, which may help assess the functional relevance of these PPIs (Figure 9).

**Fig. 9.**
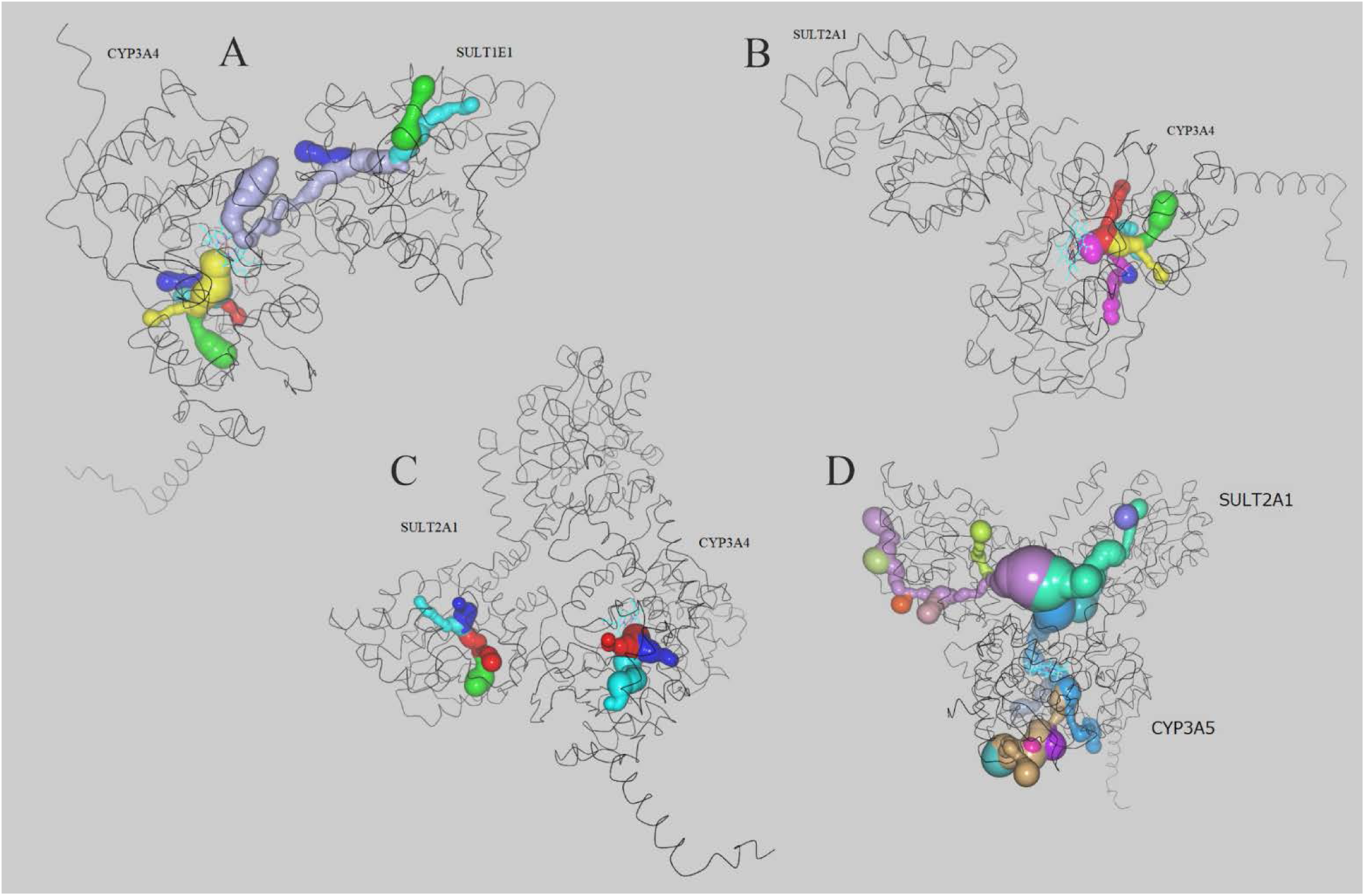
Substrate access tunnels in the modeled CYP3A4–monSULT1E1 (A), CYP3A4– monSULT2A1 (B), CYP3A4–dimSULT2A1 (C), and CYP3A5–dimSULT2A1 (D) complexes. The tunnels are colored according to the entry point on the surface of CYPs. Overlapping colors indicate merging tunnels.

## 4. Discussion

This study provides direct biophysical and functional evidence for protein–protein interactions between human CYP3A4/3A5 and SULT1E1/2A1, supporting the hypothesis that these enzymes form functional complexes. Surface plasmon resonance (SPR) assays demonstrated that these pairs form high-affinity complexes with low nanomolar dissociation constants. Binding affinity was largely independent of the PAPS cofactor, indicating that the interaction interfaces are located away from the active sites and do not block substrate access. Although the functional effects *in vitro* were modest, they exhibited distinct isoform specificity. Specifically, CYPs selectively increased SULT2A1 activity (by 38% for CYP3A4 and 108% for CYP3A5), whereas SULT1E1 activity remained unchanged. This activity enhancement may result from localized substrate channeling, where CYP3A generates hydroxylated metabolites near the SULT2A1 active site, reducing substrate dilution in the bulk phase. Alternatively, complex formation may induce mild allosteric activation. These differential regulatory effects align with the distinct physiological functions and tissue distribution of these enzymes. SULT2A1 and CYP3A4 are co-expressed primarily in the liver and small intestine, where they participate in the first-pass metabolism of steroids and xenobiotics.^50,51,64,68–71^ In contrast, SULT1E1 is a specific estrogen sulfotransferase active in steroid-sensitive tissues, such as the endometrium, prostate, and colon-where CYP3A5 is the dominant isoform.^52–60,65^ In these tissues, the CYP3A5/SULT1E1 system regulates local estrogen levels, preventing excessive hormonal signaling.^61–63,66,67^

The lack of mutual inhibition between CYP3A5 and SULT1E1 fits this physiological context, as allosteric suppression would hinder the continuous inactivation of endogenous estrogens. Thus, the CYP3A–SULT axis functions as a metabolic branch point where the physical assembly of enzymes switches the fate of common steroid substrates between oxidation and sulfation.

AlphaFold2 modeling supported these findings by predicting stable interaction interfaces (715– 1856 Å²) formed by hydrogen bonds and salt bridges. Furthermore, tunnel analysis indicated that CYP3A exit channels align with SULT entry channels, providing a theoretical structural basis for substrate channeling.^12^

However, direct extrapolation of these *in vitro* and *in silico* results to physiological conditions requires caution. First, the functional effects of the CYP–SULT interaction observed here are modest. Second, our experimental approach relies on simplified models utilizing engineered, truncated, and N-terminally modified CYP proteins. Finally, enzymatic reconstitution and interaction experiments were performed in the absence of a phospholipid membrane. Because native cytochromes P450 are membrane-bound, the lack of a lipid bilayer may alter interaction geometry, conformational dynamics, and metabolic coupling efficiency compared to the intracellular environment.

## 5. Conclusions

In summary, CYP3A and SULT isoforms physically interact *in vitro* and reciprocally modulate their enzymatic activities in an isoform-specific manner. These interactions suggest a potential dynamic mechanism for regulating the balance between oxidative and conjugative metabolic pathways. However, given the modest functional effects and the simplifications inherent to membrane-free *in vitro* models, further studies using full-length proteins in lipid-membrane systems or cellular models are required to determine the physiological relevance of these complexes.

## Abbreviations

CYPs: cytochromes P450
PAPS: 3′-phosphoadenosine-5′-phosphosulfate
pLDDT: predicted local distance difference test
PPIs: protein–protein interactions
pTMscore: predicted template modeling score
SPR: surface plasmon resonance
SULTs: cytosolic sulfotransferases

## Acknowledgments

The SPR measurements were performed using the equipment of the “Human Proteome” Core Facilities of the Institute of Biomedical Chemistry (Russia).

## Data availability

The authors declare that all the data supporting the findings of this study are contained within the paper.

## CRediT authorship contribution statement

Anastasiya Tumilovich: Conceptualization, Methodology, Investigation, Resources, Writing - Original Draft, Writing - Review & Editing, Visualization, Supervision, Project administration.

Evgeniy Yablokov: Conceptualization, Methodology, Investigation, Writing - Review & Editing, Visualization, Supervision, Project administration.

Oksana Gnedenko: Formal analysis, Investigation, Data Curation, Visualization.

Diana Mednaia: Investigation, Data Curation, Visualization.

Vadim Selin: Investigation.

Yuri Mezentsev: Investigation, Writing - Review & Editing.

Pavel Ershov: Investigation, Writing - Review & Editing.

Leonid Kaluzhskiy: Data Curation.

Tatsiana Tsybruk: Resources, Writing - Review & Editing.

Irina Grabovec: Validation, Methodology, Investigation.

Polina Shabunya: Validation, Methodology, Investigation.

Andrei Gilep: Conceptualization, Resources, Writing - Review & Editing, Supervision.

Alexis Ivanov: Funding acquisition, Resources.

All authors reviewed, edited, and approved the final version of the manuscript.

